# ZeBraInspector, a platform for the automated segmentation and analysis of body and brain volumes in whole 5 dpf zebrafish following simultaneous visualization with identical orientations

**DOI:** 10.1101/2020.10.26.353656

**Authors:** Sylvain Lempereur, Matthieu Simion, Elodie Machado, Fabrice Licata, Lilian Buzer, Isabelle Robineau, Julien Hémon, Payel Banerjee, Noémie De Crozé, Marc Léonard, Pierre Affaticati, Arnim Jenett, Hugues Talbot, Jean-Stéphane Joly

## Abstract

In recent years, the zebrafish has become a well-established laboratory model. We describe here the ZeBraInspector (ZBI) platform for high-content 3D imaging (HCI) of 5 dpf zebrafish eleuthero-embryos (EEs). This platform includes a mounting method based on 3D-printed stamps to create a grid of wells in an agarose cast, facilitating batch acquisitions with a fast-confocal laser scanning microscope. We describe reference labeling in cleared fish with a fluorescent lipophilic dye.

Based on this labeling, the ZBI software registers EE 3D images, making it possible to visualize numerous identically oriented EEs on a single screen, and to compare their morphologies and any fluorescent patterns at a glance. High-resolution 2D snapshots can be extracted. ZBI software is therefore useful for diverse high-content analyses (HCAs).

Following automated segmentation of the lipophilic dye signal, the ZBI software performs volumetric analyses on whole EEs and their nervous system white matter. Through two examples, we illustrate the power of these analyses for obtaining statistically significant results from a small number of samples: the characterization of a phenotype associated with a neurodevelopmental mutation, and of the defects caused by treatments with a toxic anti-cancer compound.

## Introduction

High-content imaging (HCI) and high-content analysis (HCA) at medium or large scales traditionally involve cell culture experiments, including the use of advanced 3D cell cultures or organoids. However, animal models remain indispensable, and small-animal models, such as invertebrates or aquatic models, are clearly more appropriate for medium or large scale analysis than larger mammalian species.

The need for HCI and HCA has greatly diversified. Various pipelines involve simple 2D imaging methods (Beaumont et al., 2015; Staal et al., 2018), 2D+time (Cornet et al., 2017), high-throughput-techniques in 2D (Jarque et al., 2018) and pseudo-3D (optical projection tomography (OPT) (Xiaoqin Tang et al., 2016). Automated systems have been developed, including some that deliver zebrafish eleuthero-embryos (EEs) in capillaries (vertebrate automated screening technology (VAST) (Early et al., 2018; Guo et al., 2017; Pulak, 2016)

For HCI, light-sheet microscopes are fast, but can analyze only a limited number of mounted samples per batch. Confocal microscopes can be used for medium-throughput imaging, but are of limited use for the imaging of deeper structures (Bruneel and Witten, 2015), due to refraction at the interface of adjacent tissue components with different refractive indices. However, tissue-clearing methods have recently progressed sufficiently for deeper imaging in a number of species, including zebrafish (Affaticati et al., 2017; Susaki et al., 2014). One of the key steps for rendering tissues optically clear is matching the refractive index (RI) of the samples to that of the mounting medium.

Zebrafish, a vertebrate model, is very similar to humans in terms of its biological and physiological properties and its body plan (Planchart et al., 2016; Roper and Tanguay, 2018), and studies in this species are subjected to fewer ethical concerns. Indeed, EEs, which do not feed, are not covered by the European directive on animal experimentation (Halder et al., 2010). This model has several advantages for HCI, including low breeding costs, high fecundity and transparency. EEs are small, and can therefore be studied in high-throughput settings. A very large collection of zebrafish mutants is now available. More than 500 are annotated in the Zfin database as having brain development defects (Davis et al., 2014). In most cases, phenotyping is performed with a dissecting microscope or the annotations are extracted from published studies.

We developed the ZeBraInspector (ZBI) platform, which uses rapid labeling and tissue-clearing protocols, dedicated 3D-printed mounting stamps and a fast-automated confocal laser scanning microscope. The use of 3D imaging overcomes the problems encountered with 2D images, which may provide misleading information about the shape of an organ or tissue if the orientation of the sample is imperfect. Registered 3D imagesalso speed up the mounting step, by making it possible to align 3D images with imperfect orientations.

This platform includes the ZBI software (https://tefor.net/portfolio/zebrainspector), which has three features. Firstly, numerous aligned samples, registered by different methods, can be visualized simultaneously on a screen, with a reference dye channel, and two fluorescence patterns in two other channels. This property is useful for many different applications, and high-resolution snapshots can be obtained in any orientation. Secondly, this software performs automatic segmentation on whole EEs and the white matter of their brains, based on staining with lipophilic dyes (such as DiI or DiO). The interface, showing numerous aligned samples, can be used to validate segmentations and to select samples for volumetric analysis. Finally, the software computes the volumetric analysis of the segmented domains.

We highlight the usefulness of the ZBI platform here, by focusing on a mutant with potential microcephaly and a long-prescribed chemotherapeutic drug found as a contaminant in the environment, with effects on aquatic animals and potential teratogenic effects in humans.

## Materials and Methods

### Zebrafish housing and husbandry

Fish were reared at a density of five adults per liter, under controlled conditions, according to European recommendations for zebrafish housing and care (Aleström et al., 2019). The environmental parameters were as follows: photoperiod = 14 h/10 h light/dark, temperature = 26.5 ± 1 °C, pH = 7.8 ± 0.1; conductivity = 240 ± 30 μS/cm, NH_4_^+^ = 0 mg/L, NO_2_^-^ = 0 mg/L, NO_3_^-^< 50 mg/L. Fish were fed on rotifers (*Brachionus plicatilis*, > 500/fish/day) for the first two weeks of feeding, and then with brine shrimps (*Artemia nauplii*, ~250/fish/day) and dry food (Skretting, Gemma Micro, to apparent satiation, two times/day). The lines used in these studies were AB (from the European Zebrafish Resource Center, Germany), casper (from the Zon laboratory, Harvard University, USA), *Tg(kdrl:DsRed)* (obtained from the Betsholtz laboratory), *Tg(fli:GFP)* and the *wdr12^hi3120Tg^* mutant line (from the Zebrafish International Resource Centre, Eugene, Oregon). All procedures were performed in accordance with European Union Directive 2011/63/EU, with the approval of the local ethics committee (no. 59 CEEA).

### Four rapid staining protocols for lipophilic carbocyanine dyes (DiI, DiO, DiD, DiR), fluorescent proteins, and optional staining with various antibodies

#### Single-dye staining (Fig. 2A)

Five days post-fertilization (dpf), EEs were killed and immersed overnight in fixative solution (4% formaldehyde in 1 x phosphate-buffered saline (PBS), 0.1% Tween 20 (PBST) at 4 °C. They were processed immediately or stored for up to several weeks at 4 °C in 0.5% formaldehyde and 0.05% sodium azide in PBST. The EEs were thoroughly washed in PBST for two days and incubated in 0.5 x saline sodium citrate buffer (SSC) with 0.1% Tween 20 for 1 h at room temperature (RT). Melanocytes were bleached by incubating the EEs in fresh depigmentation solution (0.5 x SSC, 5% formamide, 3% H_2_O_2_) for 1 h at RT, and the samples were then washed overnight in PBST. The samples were blocked by incubation for at least 5 h in blocking solution (10% normal goat serum, 10% DMSO, 5% PBS-glycine, 0.5% Triton X-100, 0.1% deoxycholate, 0.1% NP40 and 0.1% saponin in PBST) at room temperature. They were then briefly rinsed in PBST and labeled by incubation for three days with 1 μg/mL lipophilic dye (DiI, DiO, DiA, DiD or DiR, L7781, Thermo Fisher Scientific) in labeling solution (1 x PBST supplemented with 2% normal goat serum, 20% DMSO, 0.05% sodium azide, 0.2% Triton X-100, 10 μg/mL heparin and 0.1% saponin) at RT with gentle shaking. Samples were then subjected to refractive index (RI) matching and mounting.

#### Dye staining with the preservation of native fluorescent proteins (Fig. 2B-C)

Five days post-fertilization (dpf), EEs were killed and immersed overnight at 4 °C or for 2 hours at room temperature in fixative solution (4% formaldehyde in 1 x PBS, 0.1% Tween 20 (PBST)). They were processed immediately or stored for up to several weeks at 4 °C in PBST supplemented with 0.5% formaldehyde and 0.05% sodium azide. They were thoroughly washed in PBST for two days and immersed in blocking solution (10% normal goat serum, 10% DMSO, 5% PBS-glycine, 0.5% Triton X-100, 0.1% deoxycholate, 0.1% NP40 and 0.1% saponin in PBST) for at least five hours at room temperature. Samples were briefly rinsed in PBST and labeled by incubation for three days with 1 μg/mL lipophilic dye (DiI, DiO, DiA, DiD or DiR, L7781, Thermo Fisher Scientific) in labeling solution (1 x PBST supplemented with 2% normal goat serum, 20% DMSO, 0.05% sodium azide, 0.2% Triton X-100, 10 μg/mL heparin and 0.1% saponin) at RT, with gentle shaking. Samples were then subjected to refractive index (RI) matching and mounting.

#### Dye staining coupled with immunodetection (Fig.2D-H)

We proceeded as follows, to combine whole-mount lipophilic dye staining with immunodetection. Five days post-fertilization (dpf), EEs were killed and immersed overnight in fixative solution (4% formaldehyde in PBST) at 4 °C. They were processed immediately or stored for up to several weeks at 4 °C in PBST supplemented with 0.5% formaldehyde and 0.05% sodium azide. They were thoroughly washed in PBST for two days and incubated in 0.5 x saline sodium citrate buffer (SSC) supplemented with 0.1% Tween 20 for 1 h at RT. Melanocytes were then bleached by incubation in fresh depigmentation solution (0.5 x SSC, 5% formamide, 3% H_2_O_2_) for 1 h at RT, and the samples were then washed overnight in PBST. The samples were blocked by incubation in blocking solution (10% normal goat serum, 10% DMSO, 5% PBS-glycine, 0.5% Triton X-100, 0.1% deoxycholate, 0.1% NP40 and 0.1% saponin in PBST) for at least five hours at room temperature, and were then briefly rinsed in PBST and incubated with primary antibodies in staining solution (1 x PBST buffer supplemented with 2% NGS, 20% DMSO, 0.05% sodium azide, 0.2% Triton X-100, 10 μg/mL heparin and 0.1% saponin) for three days at 37 °C with gentle shaking. Samples were washed for at least 1 h in PBST at RT and incubated with secondary antibodies in staining solution without saponin for three days at 37 °C in the dark, with gentle shaking. Samples were washed for at least 2 h in PBST and stained by incubation with 1 μg/ml of lipophilic dye (DiI, DiO, DiA, DiD or DiR, L7781, Thermo Fisher Scientific) in staining solution without saponin for three days at RT with gentle shaking. The samples were then subjected to refractive index (RI) matching and mounting.

5-FU and *wdr12* samples were immunolabeled with anti-HuC/D antibodies (not shown) (ab210554, abcam, used at a dilution of 1:600). Anti-GFP (GFP-1020, Avès Laboratories) and anti-DsRed (632496, Ozyme) antibodies were used at dilutions of 1:400-1:600. The secondary antibodies used were Alexa Fluor 555-conjugated goat anti-rabbit (A-21428, Thermo Fisher Scientific, used at a dilution of 1:600), Alexa Fluor 488-conjugated goat anti-chicken (A-11039, Thermo Fisher Scientific, used at a dilution of 1:600).

### Mounting and refractive index (RI) matching

We used a RI matching medium (50% sucrose (w/v), 20% nicotinamide (w/v), 10% triethanolamine (w/v) and 0.1% Triton X-100 (v/v)), referred to hereafter as MD+, a modified form of the CUBIC-2 (Susaki et al., 2015) solution. We replaced the urea of CUBIC-2 with nicotinamide, which has a strong optical clearing ability (Tainaka et al., 2018). The RI of MD+ was adjusted, with water, to 1.457. Dye-stained samples were first incubated in 0.5 x MD+ (50% (vol/vol) in dH_2_O) for at least two hours, and were then transferred to MD+ solution. Mounting plates were coated with hot agarose/MD+ (1.5% and 2% agarose solubilized in MD+ for standard 60 mm Petri dishes and single-well plates, respectively) and appropriate stamps were rapidly positioned. The stamps were removed after the agarose hardened. Using a paintbrush, we carefully positioned the fish in the resulting niches, encasing them in small drops of phytagel/MD+ (1% phytagel solubilized in MD+) to stabilize them during subsequent steps. The mounting plates were filled with MD+ and the RI was checked several hours later. Plates with a RI matching 1.457 were imaged. If the RI did not match, the MD+ solution was renewed as often as required to reach this value.

### 3D-printed mounting molds

We used the open source CAD software Openscad (Kintel and Wolf, 2020), the files of which are hosted on gitlab:

stamp for 12 samples: https://gitlab.com/sylv.lempereur/tcfmodels/-/blob/master/petri_whole_5dpf_dr.scad
stamp for 48 samples: https://gitlab.com/sylv.lempereur/tcfmodels/-/blob/master/halfPlate_whole_5dpf_dr.scad
stamp for 96 samples: https://gitlab.com/sylv.lempereur/tcfmodels/-/blob/master/plate_whole_5dpf_dr.scad
handle for all stamps: https://gitlab.com/sylv.lempereur/tcfmodels/-/blob/master/handle.scad

These files were sliced with Cura 4 (Ultimaker, Netherlands) and printed in polylactic acid (PLA) on a standard Ultimaker 2^+^ (Ultimaker, Netherlands) with a 0.25 mm nozzle. We printed the stamps upright and standing on the straight edge of the long side, to achieve the highest resolution and fidelity in the ridges forming the small niches for the EEs. All stamps were handled with one to two default handles, which were printed in bulk separately and affixed to the stamp with superGlue (cyanoacrylate, CA).

### Imaging plates

For small batches (up to 12 EEs), we used standard 60 mm Petri dishes (Greiner Bio One International, catalog number: 628160) with the 12-sample stamp. Larger batches were mounted with the stamps for 48 or 96 specimens in single-well plates (Thermo Fisher Scientific, #140156). This system provided ample space for the objective of the microscope and had the same outer dimensions as a standard 96-well plate (Fig. 4C). This made it possible to mount the dish on the stage of a microscope with standard hardware.

### Confocal imaging

Due to the large number of samples we needed to process, our HCI system required an efficient, high-speed image acquisition setup. We therefore designed a setup based on a NIKON A1R HD microscope combined with a long-range piezo scanner objective (2 mm), a fast motorized stage and a low-magnification objective lens (10 x, 0.5 NA).

This microscope incorporates a high-definition resonant scanner, a band-pass (BP)-filtered four-channel detector unit equipped with two high-sensitivity GaAsP detectors (BP 525/50 & 595/50) and two photomultiplier tubes (PMT) (BP 450/50 & 700/75). The laser unit is equipped with four laser diodes (405 nm, 488 nm, 561 nm, 640 nm. Power > 15 mW), with switching via an acousto-optic-tunable filter (AOTF). We chose a fast motorized stage (Prior H101P2) to allow high-speed plate displacement in the *x/y* plane.

For optimization of the field of view, we combined the Nikon A1R microscope, which has a large field of view (18 mm), with a CFI Plan Apo 10XC Glyc (0.5 NA) immersion objective (Nikon), to allow the acquisition of images (tiles) with *x-y* sizes of 1200 x 1200 μm, such that the 5 dpf fish (diameter: 800 μm, length: 4 mm) could be captured in their entirety with only five tiles. This setup also provides a working distance of 5.5 mm and can correct for wide range of refractive indices, from 1.33 to 1.51.

We used a fast-piezo objective scanner (PIFOC, Physik Instrumente, Karlsruhe, Germany), which allows high-speed acquisitions over 2 mm in the *z* direction to ensure rapid image acquisition in the *z*-plane.

### Filename-based data management for HCA

We developed a data management system, using structured file names, to maintain flexibility in the choice of analytical tools and accessibility for diverse users. The data organization can be maintained even after export to a different environment, such as the laboratory of a coworker or a user of the service unit, because it is independent of a centralized tracking database.

The structured file name constitutes the ‘globally unique identifier’ (GUID) for each sample and is as easily readable to humans as it is to machines, due to its consistent structure.

For images, this file name consists of a combination of condensed composite IDs, separated by underscores: scanID_specimenID_experiment-descriptor_data-descriptor. Each of these IDs can be broken down into less complex descriptors:

- The scanID is the date of acquisition in yymmdd (year, month, day; two digits each, zero-padded) format followed by the initial of the person who acquired the image (capitalized) and an alphabetic counter (a-z) counting the scans performed by this person on this day. Having the scanID in this form and at this position forces the files into chronological order, thereby making it easier to find (potentially related) files from the same session in a file management system (Explorer, Finder, Nautilus).
- The specimenID consists of the experimentID, which identifies a batch of samples from a project, and an alphabetical counter, identifying each specimen individually.
- The experiment-descriptor contains a shorthand descriptor of the developmental stage of the specimen, the projectID it belongs to, an X-voxel count, providing information about the resolution of the image, a freeform comment for non-HCS-acquisitions and a machineID, which identifies the microscope and the technique used to acquire the image. The projectID can also be broken down into the abbreviated project name, a numerical counter and an abbreviation of the Latin species name of the specimen, all separated by hyphens.
- The data-descriptor differentiates between the different stages of processing and analysis of the image. As all the data derived from a given data set carry the same base name (the first three descriptors), the data-descriptor informs the user of the status for the mode of data considered.

Under individual-image conditions, file names are constructed interactively with a helper application that checks the integrity of the GUID. However, names cannot be assigned directly by the microscope software, and we therefore developed a software tool allowing users to re-establish the association between the experimental history of the sample and its HCI image stack by translating the file name assigned by the microscope into the appropriate file name for the ZeBraInspector system.

This process is based on a pseudodatabase system consisting of two spreadsheets (one for input, the other for output) and a couple of javascript and python scripts. These scripts ensure that the function of the filename as a GUID is not damaged, by checking that the right file name components are applied to the correct image and preventing duplications. The correlation between the image and the underlying sample is thus established on the basis of the position of the sample in the mounting grid. This information is conveyed in the microscope-generated file name and entered into the input spreadsheet. Careful user guidance on the input spreadsheet facilitates the use of this spreadsheed and increases user confidence.

### *wdr12* mutants

*wdr12^h^*i3120Tg mutants were obtained from the Zebrafish International Resource Center (ZIRC, Eugene, OR, USA). Adult zebrafish were maintained as heterozygotes and inbred to generate homozygous mutant embryos. EEs were sorted under a stereomicroscope (Olympus SZX16) on day 5. Almost 25% of these EEs had eyes that appeared to be smaller than those of non-mutant EEs at the same developmental stage and were considered, *a priori*, to be mutants. Putative mutants and their siblings were processed according to the zFaCT protocol. After imaging, all fish were unmounted and subjected *a posteriori* to genotyping (not shown). Wildtype and mutant *wdr12* alleles were detected with the following primers: forward 5’-ACCCAGCTGACATTTGCTCT-3’, reverse 5’-TTCTTGGCTTCCAGCAGTTT-3’ and reverse2 5’-CAATATCACCAGCTGAAGCCTA-3’. PCR with the forward and reverse primers yielded a 244 bp amplicon for the wild-type allele, and an 831 bp amplicon for the mutant allele.

### 5-FU treatments

5-fluorouracil (Sigma, USA) was applied in DMSO, which was used as a carrier. Three types of control conditions were used: zebrafish in embryo medium, zebrafish placed in six-well plates in 5-FU-containing medium, zebrafish in embryo medium+DMSO. The 5-FU application protocol is described in the results section. At 5 dpf, all EEs were imaged under the dissecting microscope. The EEs were then mounted in 48-well molds to ensure homogeneous acquisitions between samples with our fast confocal microscope. The microscopy parameters were kept constant between the three successive experiments.

### Image processing and statistical analysis

We performed registration, segmentation, statistical analysis and graph plotting in a python 3.7 environment under Ubuntu 16.04 and Windows 10, on an HP computer with an Intel Core i7 vPro CPU and 32 GB of RAM. We used Numpy, Scipy (Virtanen et al., 2020), scikit Image (van der Walt et al., 2014; Yaniv et al., 2018), matplotlib (Hunter, 2007), openpyxl, Pink (Couprie, 2011) and SimpleITK (Yaniv et al., 2018).

Statistical analysis was performed as follows: Fisher’s exact test was performed to compare the variances of two clusters. If the variances were found to be equal, Student’s *t* test was performed. Otherwise, a Welch test was performed.

We manually performed 3D segmentations of whole EE and their white matter, using Amira for Life & Biomedical Sciences software (Thermo Fisher Scientific, USAWe then compared these segmentations with those obtained automatically.

We assessed segmentation accuracy, by calculating Matthews’ correlation coefficient (MCC), the Sørensen-Dice coefficient (DSC) and a general balanced metric (Cappabianco et al., 2019) setting ***δ*** = 2 (GB2). DSC and GB2 did not take true negatives into account. This problem was counterbalanced by computation of the MCC. However, the large numbers of black voxels around the sample could create computational artifacts. We prevented these artifacts by calculating MCC in a bounding box around the union of the manual and automatic segmentations, with a tolerance of 5 voxels. We also calculated GB2 because this coefficient favors false positives over false negatives. When combined with DSC and MCC, this led to a better estimation of segmentation accuracy.

Our ZeBraInspector (ZBI) graphical interface was developed with the Pyside2 library. The workspace screen displays up to 6×3 views simultaneously. Rotations, slice stacking, section moves and contrast enhancement are computed in real time with the Numba library to obtain Python and Numpy code acceleration. The software was thus built exclusively from native Python code.

## Results and discussion

We established the ZeBraInspector (ZBI) platform for high-content imaging (HCI) and high-content analysis (HCA) of 3D images for studies of zebrafish neurodevelopment. Based on the observation that lipophilic tracers yield intense labeling in the brain, we designed a simple standardized procedure for staining, mounting and analyzing whole 5 dpf EEs with these dyes. This procedure is currently tested and adapted with other samples: it works in our hand with 3dpf fish (eye-less registration mode) and up to seven days (not shown). ZBI may certainly be used with other fish species such as *Astyanax* or medaka.

Staining made it possible to register signals, resulting in a gain of time at the mounting step by allowing the use of poorly oriented samples. The software also performs volumetric estimations of the whole body and of the brain matter. All steps have been optimized to minimize time and effort and to increase the generalizability of the experiment. For a batch of 48 samples, only about eight days are required for completion, from initial fixation to the quantification of brain and whole EE volumes (Fig. 1).

**Fig. 1.**
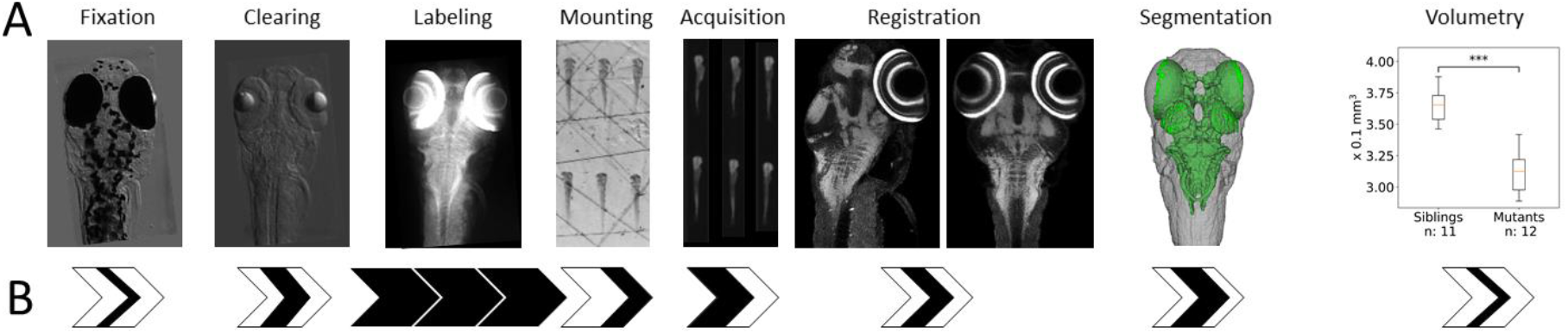
The ZeBraInspector (ZBI) platform. (A) The ZeBraInspector platform involves several steps. After fixation, a tissue clearing process is performed to reduce light diffusion and absorption, and the samples are then stained with a lipophilic dye. Samples are mounted in specific molds for rapid automatic acquisition on a microscope. The 3D images can be registered or segmented, and a volumetric analysis of the EE and white matter can be performed. (B) Time required for each step of the ZeBraInspector platform. Each arrow represents a day of treatment. Black sections correspond approximately to the time required for one day of experimentation. The whole process takes about 10 days from fixation to volumetric analysis.

### High-content 3D image analysis requires reliable features for automatic segmentation: a robust labeling and tissue clearing protocol

High-content analysis (HCA) requires consistent fluorescence signals for automatic segmentation. We tested lipophilic dyes of the carbocyanine family (e.g. DiO, DiI and DiD) for reliable visualization of the main morphological features of the EEs (Fig. 2 and Fig. 3). When applied as a counterstain, which is facilitated by the small size of the EE, this type of dye, which is traditionally used for electrode marking and neuronal tracing, strongly stains the membrane-rich white matter of the brain. Due to the lower density of membranes in non-neural tissues, the staining was less pronounced in these tissues than in the brain, but nevertheless sufficient for automatic segmentation of the whole body. We established a staining protocol yielding optimal permeabilization while preserving the shape of the EE.

**Fig. 2:**
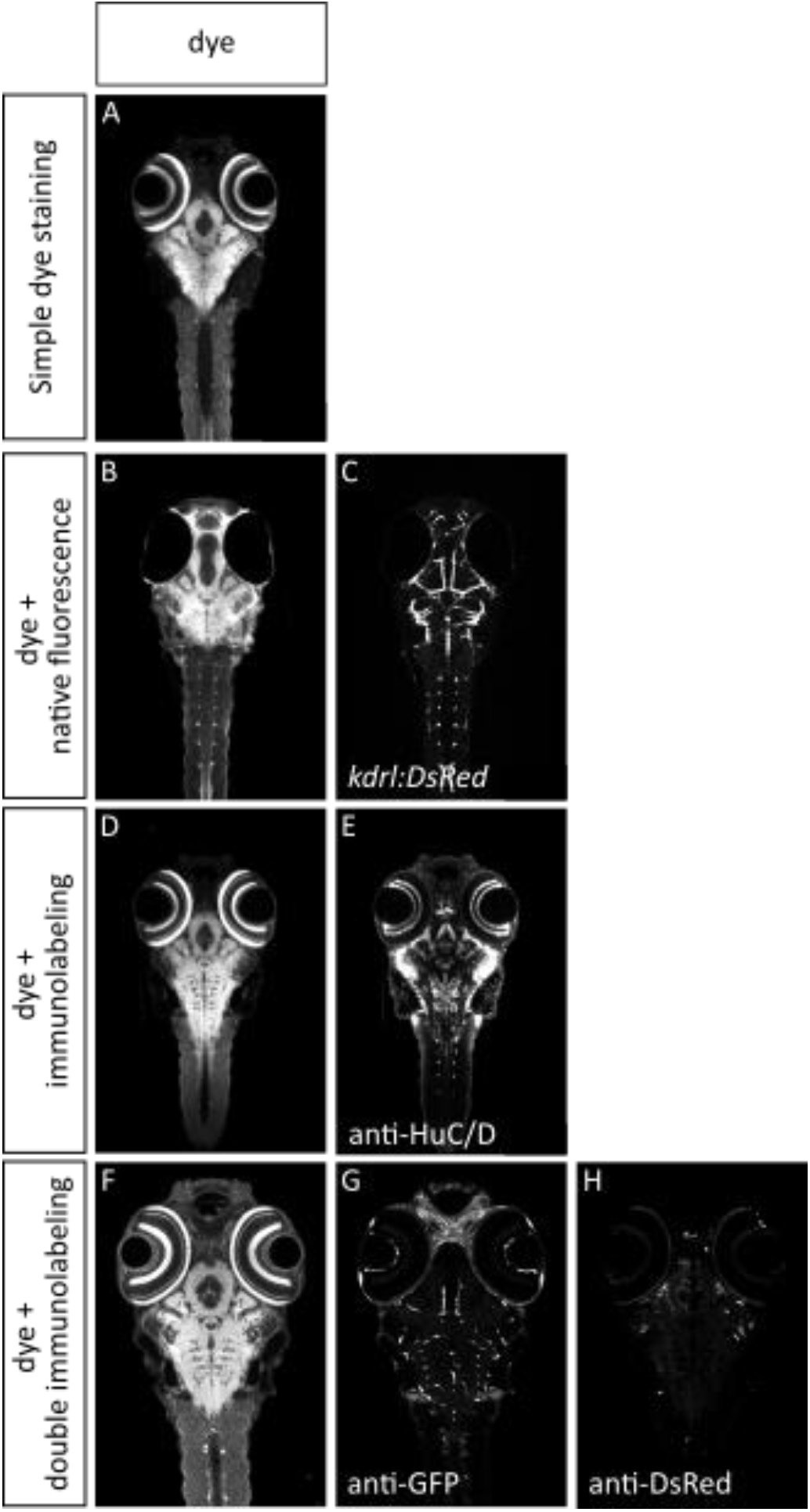
Four examples of fast staining protocols compatible with the ZeBraInspector (ZBI) platform. (A) Confocal horizontal optical section of a 5 dpf zebrafish after DiO staining. (B-C) The zFaCT labeling protocol is compatible with native fluorescence imaging. A 5 dpf EE from the *Tg(kdrl:DsRed);casper* transgenic reporter line, in which the endothelial cells of the blood vasculature are counterstained with DiO. (B) Confocal horizontal optical section of the DiO channel. (C) The same optical section, showing DsRed fluorescence. (D-E) The zFaCT labeling protocol allows whole-mount immunolabeling. A 5 dpf EE incubated with an anti-HuC/D antibody to label neurons and counterstained with DiO. DiO and anti-HuC/D antibody staining patterns from the same horizontal optical section are shown in D and E, respectively. (F-H) Three-channel imaging was performed on immunolabeled 5 dpf EE from the *Tg(fli:GFP)* transgenic line into which SINV-mCherry/2A was injected on day 3. DiD, GFP (anti-GFP antibody) and mCherry (anti-DsRed antibody) fluorescence patterns from the same horizontal optical section are shown in F, G and H, respectively.

**Fig. 3.**
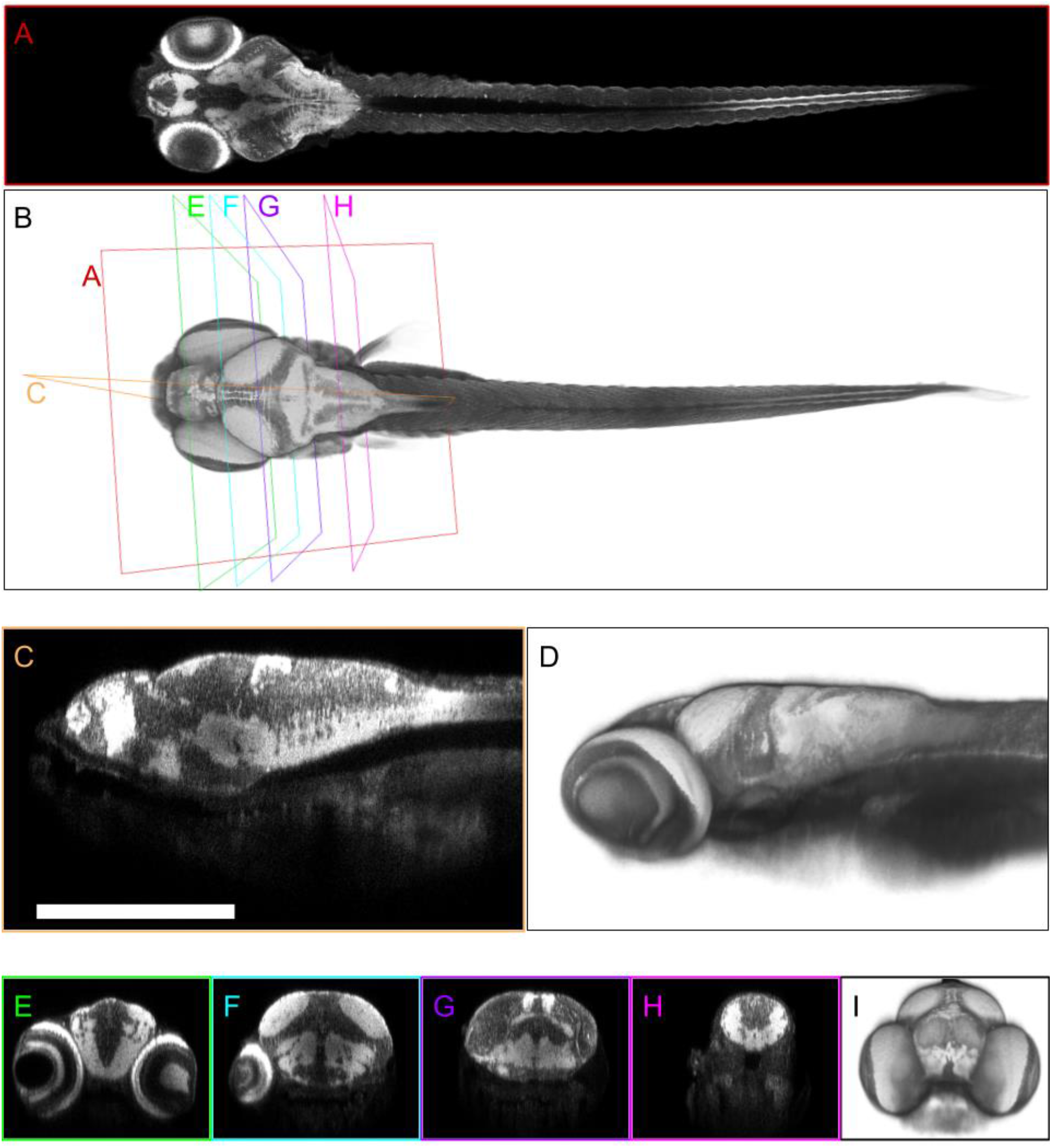
The lipophilic dye DiO: a reliable marker visualizing brain white matter and whole EE. Zebrafish EE (5 dpf) were labeled with the lipophilic marker DiO and cleared. (A, C, E-H) Optical sections, 2 microns thick (raw data), from confocal microscopy are shown in Fig. 4. Section planes are shown in B. B, D, I: 3D volume rendering of the EE (raw data). (A) Horizontal section, imaging plane. (B) Dorsal view. (C) Sagittal section (posterior region cropped). (D) Lateral view of the anterior region of the EE. (E-H) Transverse sections. (I) Frontal view of the EE head.

This protocol can be tuned for direct recording of the native fluorescence of reporter lines (Fig. 2B-C), skipping the depigmentation step, provided that an unpigmented genetic background is used (Materials and Methods). This approach is scalable with an optional immunolabeling step using anti-GFP/RFP or other antibodies (Fig. 2D-H) if additional morphological data are needed.

Sample clearing is necessary for confocal microscopy, because a slight signal loss would otherwise be observed with increasing depth in 5 dpf zebrafish. Indeed, homogeneously contrasted images are required for the subsequent automatic segmentation steps, and along the *z* axis of the image stack. Incubation in mild detergents leads to a partial delipidification, promoting sample clearing. The imaging buffer is crucial for successful tissue clearing, as it renders the refractive index (RI) of the sample equal to that of the surrounding medium, thereby rendering cell membranes, cytoplasmic proteins and other cell components transparent. After testing several products and concentrations, we opted for a medium based principally on sucrose, nicotinamide and triethanolamine (MD+, see the Materials and Methods section), which has suitable optical parameters for imaging with dipping lens objectives with high or tunable RIs.

The DiO labeling of a representative 5 dpf zebrafish is shown in Fig. 3. As shown in Fig. 3A, DiO labeling is observed throughout the body, but is strongest in areas in which the density of myelinated fibers is high. Retina photoreceptors, olfactory bulbs and the main fiber tracts of the brain display a very strong DiO signal, making it possible to detect fine anatomical structures and landmarks (Fig. 3 E-H: telencephalon, optic commissures, lateral forebrain bundle, optic tectum, medial longitudinal fascicle). Moreover, all anatomical structures are labeled. Outside the brain, ganglions and motor neurons (Fig. 3A), appearing as dots along the spinal cord, are strongly positive. Muscles (Fig. 3A) and other non-neuronal structures (Fig. 3C: yolk, digestive tract and organs, etc.) have a much weaker DiO fluorescence signal than brain structures, but nevertheless remain detectable. Overall, these data show that DiO can be used as a marker of brain structures and for the detection of whole EEs, on the basis of background labeling.

### HCI: description of rapid-mounting setups for EEs

Many mounting strategies for the high-throughput microscopy of zebrafish EEs have been published (Alessandri et al., 2017; Donoughe et al., 2018; Kleinhans and Lecaudey, 2019; Staal et al., 2018; Westhoff et al., 2013; Wittbrodt et al., 2014), but most are suitable for embryos but not for 5 dpf EEs. We therefore designed 3D printed fixtures for the generation of regular grids of small niches of approximately the shape and size of 5 dpf EEs (Fig. 4). Once placed in these narrow niches with fine paintbrushes, the EEs remain in the chosen orientation, dorsal side-up in this pipeline. This mounting strategy is possible because EEs dry out more slowly in high RI medium than in water. EEs are immobilized with a drop of phytagel, a transparent plant culture medium (Zhou et al., 2019), which serendipitously proved to be ideal for microscopy. One of the advantages of this platform is that the orientation of the EEs does not need to be perfect, provided that there is no eye in too dorsal position to disturb the acquisition of the brain region below the eye. The imaging plates and stamps are chosen according to the number of samples.

**Fig. 4.**
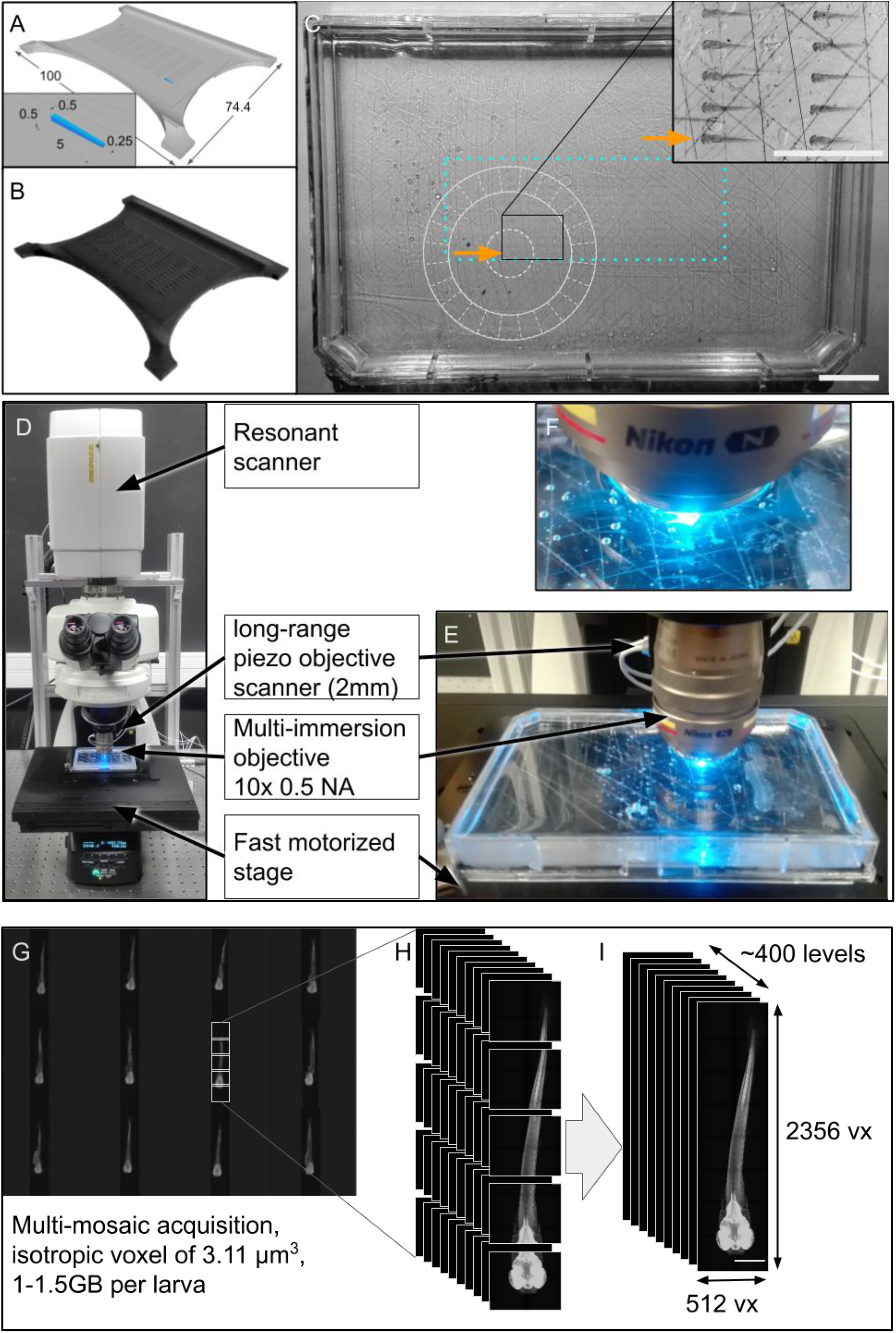
HCI tools and strategy. (A) Modeling the stamp in CAD software. All measurements are in millimeters. (B) 3D-printed plastic stamp. (C) Dissecting microscope view of a single-well plate with 48 embedded samples. The round, light-gray line diagram represents the microscope objective; this demonstrates why so much space is required between the rim of the single-well plate and the specimens. The cyan dotted rectangle outlines the overall scan area. The gray lines are scratches made manually in the plastic of the plate to strengthen the adhesion of the agarose to the dish. Scale bar: 5 mm. (D) Overview of the CLSM setup. (E) Focus on the dipping lens and the long-range piezo objective scanner. (F) High-magnification view of the imaging setup displaying the grid of embedded fish (21 dpf) (G) Partial screenshot from the Nikon acquisition software (NIS-element) displaying a multi-mosaic acquisition for 12 samples. (H) Mosaic acquisition: as the specimen is larger than the field of view, multiple tile-stacks with a 10% overlap are acquired independently. These data are recorded with a sampling rate of 512X512 voxels (vx) of 2.5 x 2.5 μm. With a correspondingly adjusted *z*-step size, this results in isotropic voxels. (I) Dimensions of a standard data set. The resulting image-stack after merging of the tile-stacks contains about 400 slices, each 512 voxels wide and 2256 voxels long.

### HCI requires speed: a description of the main advantages of fast confocal laser scanning microscopy

Light-sheet microscopy is the most powerful method for live applications, but we chose not to use it here, because it does not allow large-scale mounting and requires a high level of expertise in optics an computing (Albert-Smet et al., 2019).

We chose a commercially available fast confocal laser scanning microscope, which was reasonably priced and easy to use. The images generated were of a reasonable size. Depth-dependent signal loss is a drawback of confocal microscopy, but can be overcome by tissue-clearing techniques. Several options make rapid acquisition possible: a piezo-electric objective scanning nanofocusing system allowing high-speed movements in the *z* plane over a range of 2 mm, a large field objective allowing the imaging of an entire 5 dpf EE with only five tiles, and a high-definition resonant scanner coupled to two high-sensitivity photomultipliers.

### HCA of whole EEs and their brain white matter

#### Presentation of the pipeline

For analysis of the large amounts of data acquired, we developed computer software to automate the analysis as much as possible (Fig. 5). The first step in the analysis involves the use of a renaming tool to give the files names that are informative for the user. Robust image traceability is, indeed, essential, with links to sample names and experimental procedures.

**Fig. 5.**
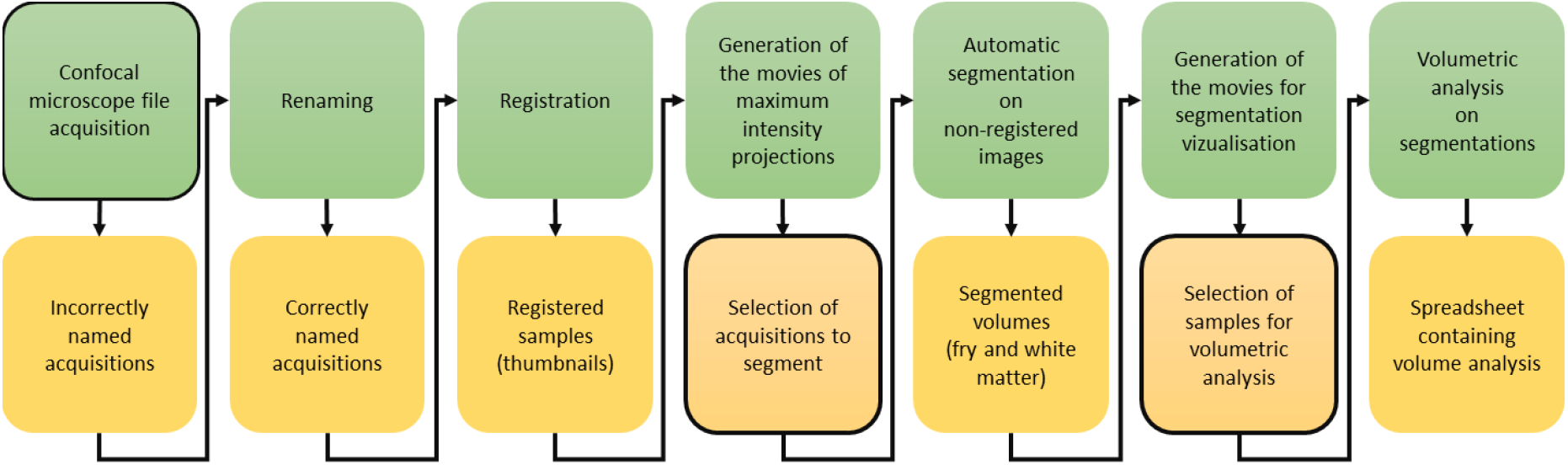
The ZeBraInspector informatic pipeline, making it possible to register, segment and visualize high-content 3D images. The ZBI pipeline consists of a succession of computation steps. The datasets acquired are renamed, registered and segmented. Segmentations are used for volumetric analysis. Framed squares represent sequences requiring manipulation by the user.

Within the ZeBraInspector (ZBI) software, registration makes it possible to simplify comparisons between samples. Maximum intensity projections of substacks of the image can be generated. This makes it possible to remove invalid samples before segmentation to reduce the computing time. These segmentations automatically detect the positions of the EE and its white matter in the original acquisition. However, segmentation errors are possible, and a new visualization is provided to the user, to make it possible to eliminate such errors. This visualization displays the segmentations for comparison with the raw data. Once any incorrect segmentations have been removed, ZBI performs a volumetric analysis.

#### Registration for faster batch visualization of EEs

Comparisons between samples were facilitated by implementing several procedures for sample registration in ZBI. We provide a detailed description below of the first of these procedures, using the lenses as landmarks.

The original image (Fig. 6A,E) is morphologically closed, and then morphologically opened to smooth the signal. This smoothed image is then thresholded to keep the top ten percent of gray values. A rough segmentation of the white matter is then obtained by keeping the largest element resulting from this thresholding (Fig. 6B,F). A vector describing the main orientation of the white matter is defined, retaining the largest half axis (major semi-axis) of the ellipse encompassing the thresholded voxels. The computing time required for subsequent steps is reduced by downsampling the original image to a quarter of its original size in each dimension. A rough segmentation of the whole larva is computed by thresholding the downsampled image, keeping the largest element resulting from this thresholding and then closing the image and removing internal holes. For the detection of black elements in the specimen, a watershed is computed using the lowest five per cent of gray values as interior markers, then the outside of the larva and the top 50% of gray values as external markers, and a morphological gradient of the closed image as the topological map. Lenses are detected among the dark elements by retaining the two largest spherical elements (Fig. 6 B,F). Using these two landmarks, it is, therefore, possible to obtain two vectors: the interocular vector, from the center of one crystalline lens to the other, and the translation vector, pointing from the center of the lenses to the center of the whole image. Orthogonality is ensured by projecting the vector describing the main axis of the white matter onto the plane defined by the interocular vector. A final vector is obtained by calculating the product of the interocular and projected vectors. This fourth vector is oriented to point to the part of the image with the highest mean gray value. The four vectors (Fig. 6 C, G) are then used to calculate the transformation matrix used to align the image with its axis (Fig. 6 D, H). The calculation of the translation and rotation matrices based on the DiO channel data makes it possible to apply this image registration to all signals acquired in parallel to the DiO reference channel.

**Fig. 6.**
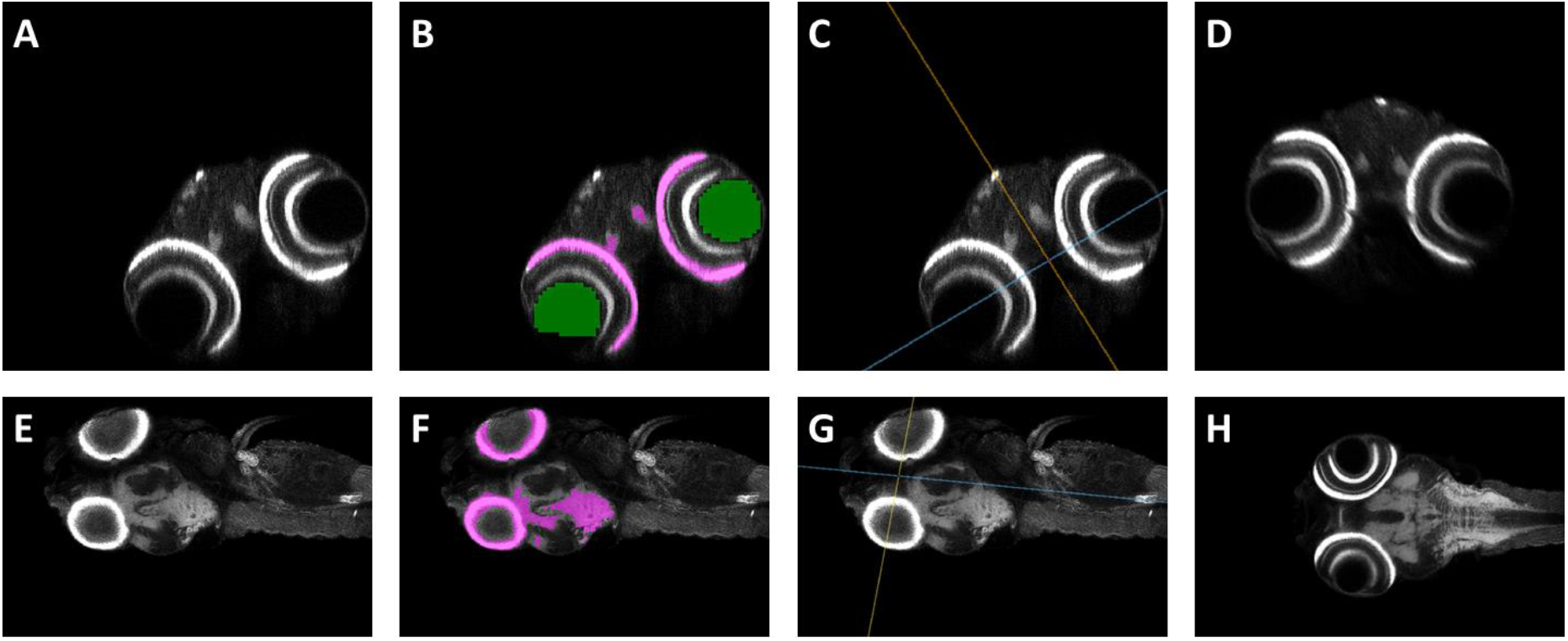
A registration method using the lenses as landmarks and the fitting of an ellipsoid to the brain white matter for EE axis definition. (A-D) Midbrain dorsal sections. (E-H) Midbrain transverse section (A,E) Raw data. (B, F) Superimpositions of segmentations of the white matter (magenta) and crystalline lenses (green) onto the raw data (gray). These landmarks are used to compute the direction of the white matter. (C, G) Superimposition of orientation planes onto raw data (gray). Yellow, blue and red lines represent the lens-to-lens, main axis and third axis planes, respectively. (D,H) Results of the registration. The sample is centered and the white matter is aligned with the main axis of the image.

However, it may not be possible to use the lenses as landmarks, if the eyes are pigmented and the light cannot penetrate them, for example. Two other registration processes, similar to that described above, have been developed to get around this problem, but are not described in detail here. For the registration of fish with pigmented eyes, the algorithm uses whole eyes as landmarks, with differences only for some morphological kernels and threshold values. A third registration does not use the eyes at all, instead computing the bounding ellipsoid from the white matter, using its axes as vectors for alignment. This option may orient samples less precisely, but we recommend its use in cases in which registration based on the lenses or eyes is unsuccessful. All the algorithms are available from the gitlab repository.

### Segmentation of the whole EE

One of the key tasks in our processing pipeline is the segmentation of the whole EE. For automatic segmentation that flexibly adapts to images displaying some diversity in terms of intensity and quality, many steps are required, regardless of the standardization efforts made. A morphological closing is initially required to eliminate local undersegmentation due to local gradients. This is followed by a morphological opening eliminating background noise external to the sample. The resulting smoothed image is first used to calculate a morphological gradient reflecting the transition of grayscale values over the image. The pixels are white in areas of abrupt transition, and dark in homogeneous regions. A topological map is obtained as an intermediate result. Two thresholds based on the gray levels of the smoothed image are then applied: the first threshold corresponds to the internal zones (body) of the sample, with high values retained, and the second corresponds to the external zones (background), with low values retained. Using these thresholds, we labeled the local maxima of the smoothed image by assigning values of 0, 1 or 2, depending on the area in which they were located. 0, 1 and 2 correspond to neutral, external and internal seeds, respectively.

The next step is a segmentation based on the watershed method, which requires seeds and a topological map. The seeds, corresponding to the values 1 or 2 defined above, are used to initialize the watershed. During the watershed procedure, these seeds grow to fill the area of interest. The algorithm is first applied to a downsampled version of the image, to locate the sample. This makes it possible limit the computation of the sample watershed when the high-resolution image is processed, thereby reducing the calculation time. However, due to residual light diffusion into tissues, the acquisition side is brighter than the opposite side. A specific threshold is calculated to remove low grayscale values on the acquisition side, to solve this problem and to improve segmentation accuracy. Finally, a morphological closure is applied to eliminate potential non-segmented areas within the EE.

For validation of this segmentation method, we imaged 12 samples with the corresponding grid and computed segmentations with our pipeline (Fig. 7A). Following segmentation, the results were checked with ZBI, and this led to two very badly labeled samples being discarded that would otherwise undoubtedly yielded abnormal segmentations. We compared the automatic segmentations (Fig. 7B-E) with those obtained manually (Fig. 7F-I). We obtained median Sorensen-Dice coefficient, general balanced metric (GB2) and Matthew’s correlation coefficient (MCC) values of about 0.95 (Fig. 7J). These high scores indicate a high degree of correspondence between the manual and automatic segmentations. On average, it took six minutes to calculate an automatic segmentation, which is not significantly different to the eight minutes required for acquisition. This calculation time, estimated on standard laptops, could certainly be reduced by parallel calculations on dedicated computation servers. In conclusion, our automatic segmentation method provides a precise segmentation of the whole EE.

**Fig. 7.**
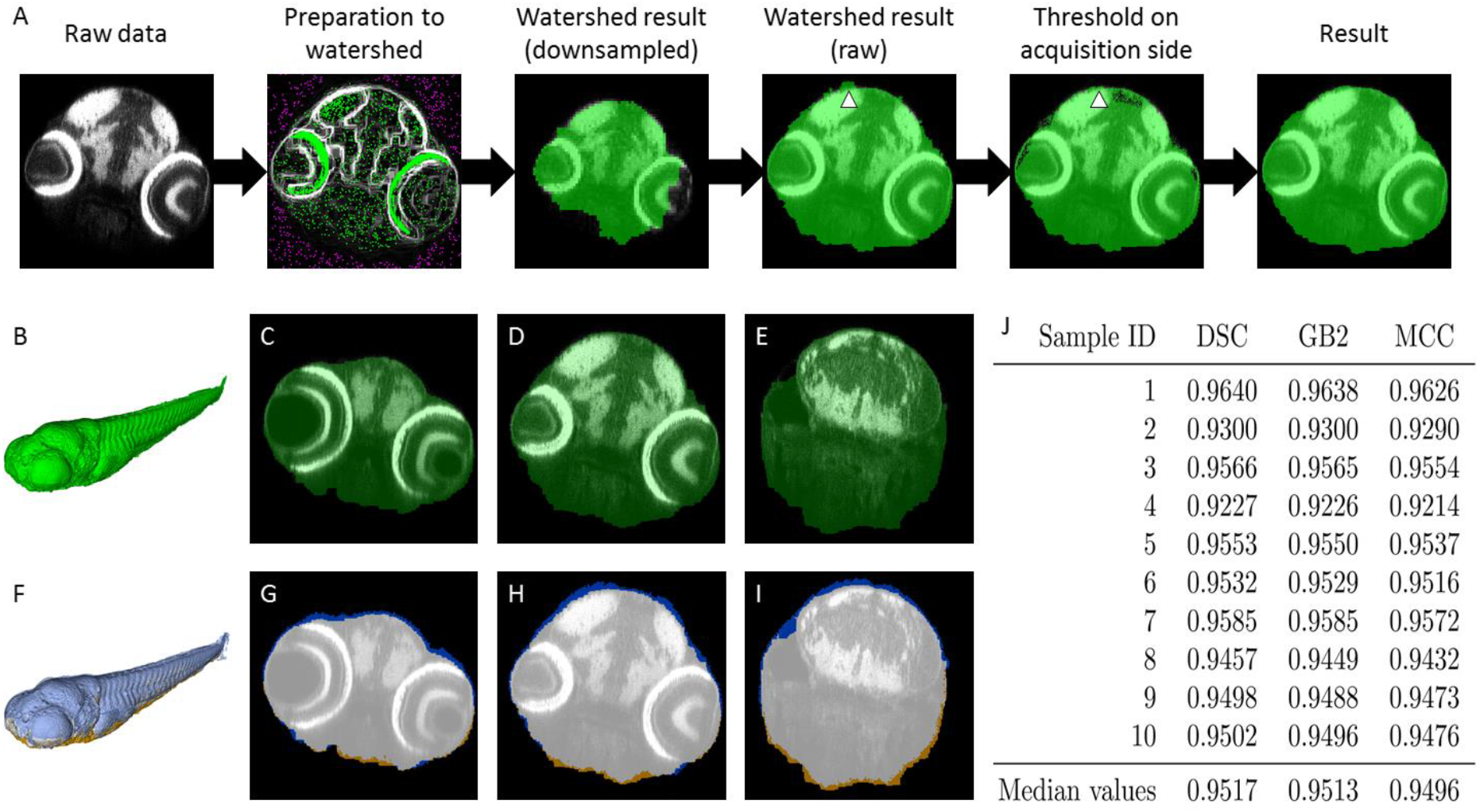
Pipeline and validation of EE segmentation. (A) Main steps in the computation of EE segmentation by watershedding. Each step is illustrated with a transverse section of the midbrain. Due to light scattering, the acquisition side is brighter than the opposite size. The watershed oversegments dorsally (arrowheads). This problem is solved by applying a specific threshold on the acquisition side. The algorithms are available from gitlab (see the Materials and Methods). (B-E) Segmentation results. Segmentation in green, raw data in gray. B. 3D surface rendering. C-E. Transverse section. (F-I) Comparison of manual and automatic segmentations. Raw data are shown in gray. Manual segmentation only, automatic segmentation only and the intersection between these intersections are shown in brown, blue and white, respectively. F. 3D surface rendering. (G-I) Transverse section. (J) MCC, GB2 and SDC coefficients for automatic segmentations compared with those for manual segmentations. The median values of these coefficients are about 0.95, indicating that the two segmentation strategies yield almost identical results.

#### Brain white matter segmentations

We made use of the ability of DiO to label brain white matter strongly, for the automatic segmentation of this tissue. As for the segmentation of the whole EE, the first step was the calculation of a morphological opening followed by a closing. These steps generate a smoothed image, which was used to calculate a first threshold for retaining the 10% highest grayscale values in the image. This thresholding creates a class of voxels that we consider to correspond to white matter.

A second thresholding is then performed on the smoothed image, to retain the lowest 50% of the grayscale values of the image. We improved the precision of the segmentation by watershedding, by eroding the thresholding results to ensure the presence of a neutral range between the two zones. As described above, we then retained the local maxima of the smoothed images within each zone to create an image including the necessary seeds for watershedding.

The presence of elements strongly labeled with DiO outside the brain, within the digestive tract for example, made it necessary to introduce a second step to refine the segmentation, limiting it to the brain. For this, we select the largest segmented element, corresponding to the brain. We then calculate a morphological expansion of this object, to create the root allowing the calculation of a geodesic expansion, using the watershedding result as a mask. This step removes elements that are not part of the brain, while retaining small isolated brain structures.

Using a strategy similar to that described above to validate the segmentation, we segmented the brain white matter of the 10 previous EEs (Fig. 8). Comparisons between automatic and manual segmentations (Fig. 8F-I) yielded median DSC, GB2, and MCC values of about 0.88 (Fig. 8J). Automatic segmentation took an mean of three minutes longer. Thus, whole-body and brain white matter segmentations took a time similar to that required for image acquisition.

**Fig. 8.**
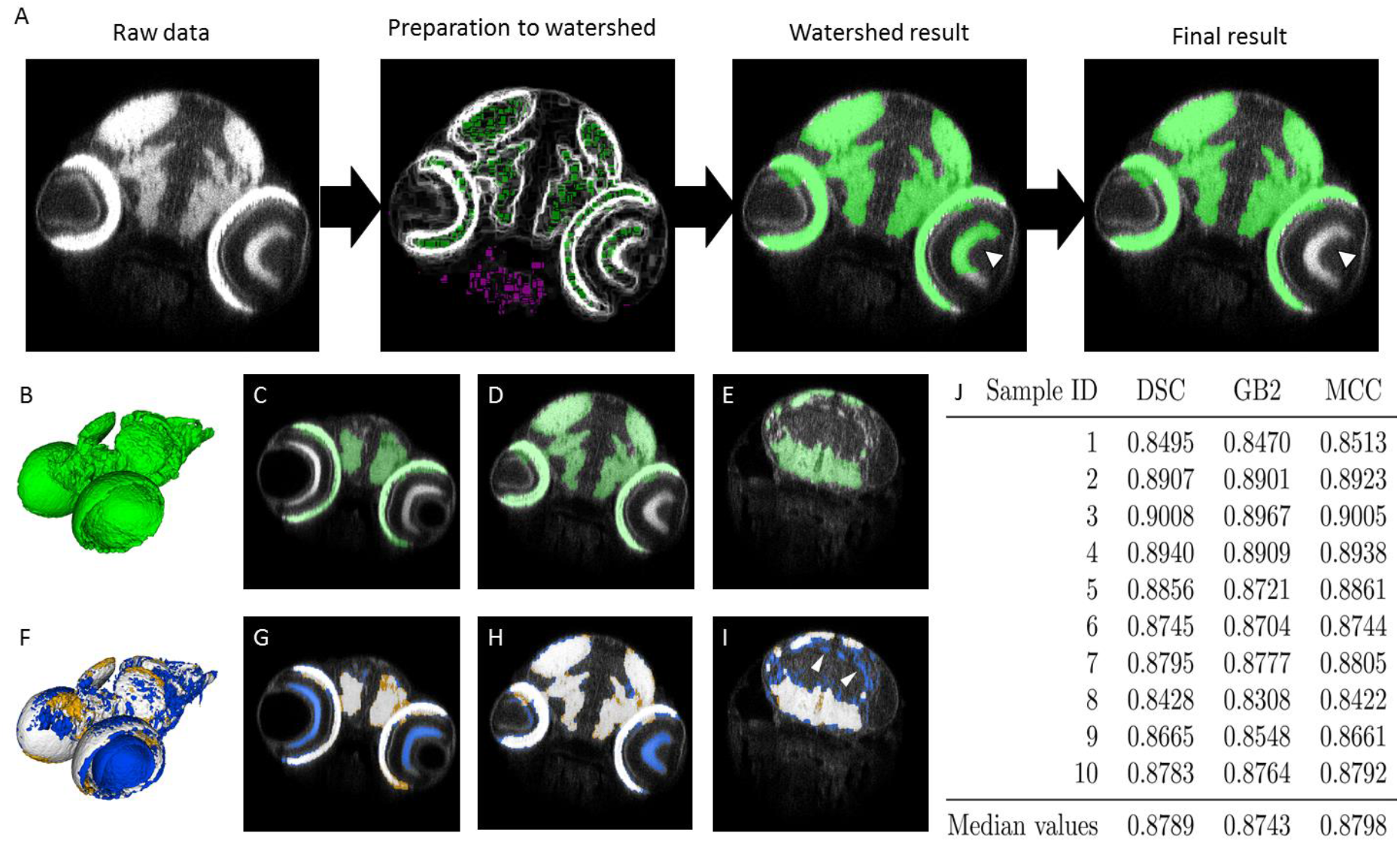
Pipeline and validation of white matter segmentation. (A) The main steps in the computation for white matter segmentation by watershedding. Each step is illustrated with a transverse section of the midbrain. Watershed computation detects unwanted elements, such as the retina (arrowhead) or yolk (not shown). These oversegmentations are removed by selecting the largest elements and their near neighbors. Details of this process are available from gitlab (see the Materials and Methods). (B-E) Results of segmentation. Segmentation in green, raw data in gray. (B) 3D surface rendering. (C-E) Transverse section. (F-I) Comparison of manual and automatic segmentations. Raw data in gray. Manual segmentation only, automatic segmentation only and intersection between the two segmentations, in brown, blue and white, respectively. (F) 3D surface rendering. (G-I) Transverse section. (J) MCC, GB2 and SDC coefficients for automatic segmentation relative to manual segmentations. Coefficients are greater than 0.8, indicating that automatic segmentations are precise enough for volumetric analysis. Indeed, segmentation errors were similar for all samples. For example, small white matter tracts in the brain are not segmented (arrowheads in I).

#### Checking of segmentation accuracy by the user

The software allows users to check segmentation quality manually, by producing an image superimposing the segmentation of the white matter on the segmentation of the whole EE. These segmentations can then be compared with raw data via ITK-SNAP or any other viewer allowing navigation within 3D images. Furthermore, this superimposition of segmentations makes possible the automatic creation of images showing the overlay of the segmentation on the raw data, on transverse sections, for example. These images can be used to detect potential segmentation errors, making it possible to remove some samples before starting statistical analyses.

Our software can also display segmentation on the desired channel, making it possible to detect segmentation failures. The software then proposes the elimination of these failures before statistical analyses.

#### Volumetric analysis

Volumetric analyses are performed by multiplying the number of segmented voxels in each segmented specimen by the physical size of the voxel, to obtain its real-world volume. This computation, when performed for both segmentations, provides two different volumes: the whole EE volume and the volume of its white matter. Dividing the volume of the white matter by the volume of the EE, we obtain a ratio providing information about more subtle defects of neural development such as impaired neurogenesis. The values obtained are automatically recorded in an Excel spreadsheet and provided to the user.

### HCA by the ZeBraInspector (ZBI) platform reveals discreet, but statistically significant brain defects

We describe here the use of our pipeline to refine the characterization of a microphthalmic mutation in a ribosome biogenesis gene. We selected the *Wdr12* mutant line, in which visible eye microphthalmia was on lateral transmission light microscopy pictures (2 and 6 dpf) was reported in the Zfin database (Fig. 10A). We crossed heterozygous founders and selected approximately one quarter of the 5 dpf EEs on the basis of microphthalmia criteria. We first confirmed on one batch of EEs that the affected EEs were homozygous for the mutation (data not shown). Eye phenotypes were 100% penetrant in *wdr12* homozygous (Hmz) embryos, whereas heterozygous (Htz) embryos were indistinguishable from their wild-type (Wt) siblings.

We assessed whether brain hypomorphy, which was not obvious on observation under a dissecting microscope, could be characterized by ZBI. We first used the snapshot button of ZBI to extract maximum-intensity projections of transverse thick sections of the midbrain (Fig. 9, see te Methods for details of the protocol and guidelines). This operation was greatly accelerated by navigation in parallel in registered samples, visualized in the same orientation and at identical levels. ZBI can thus be used for efficient rapid analyses of images in batches.

**Fig. 9.**
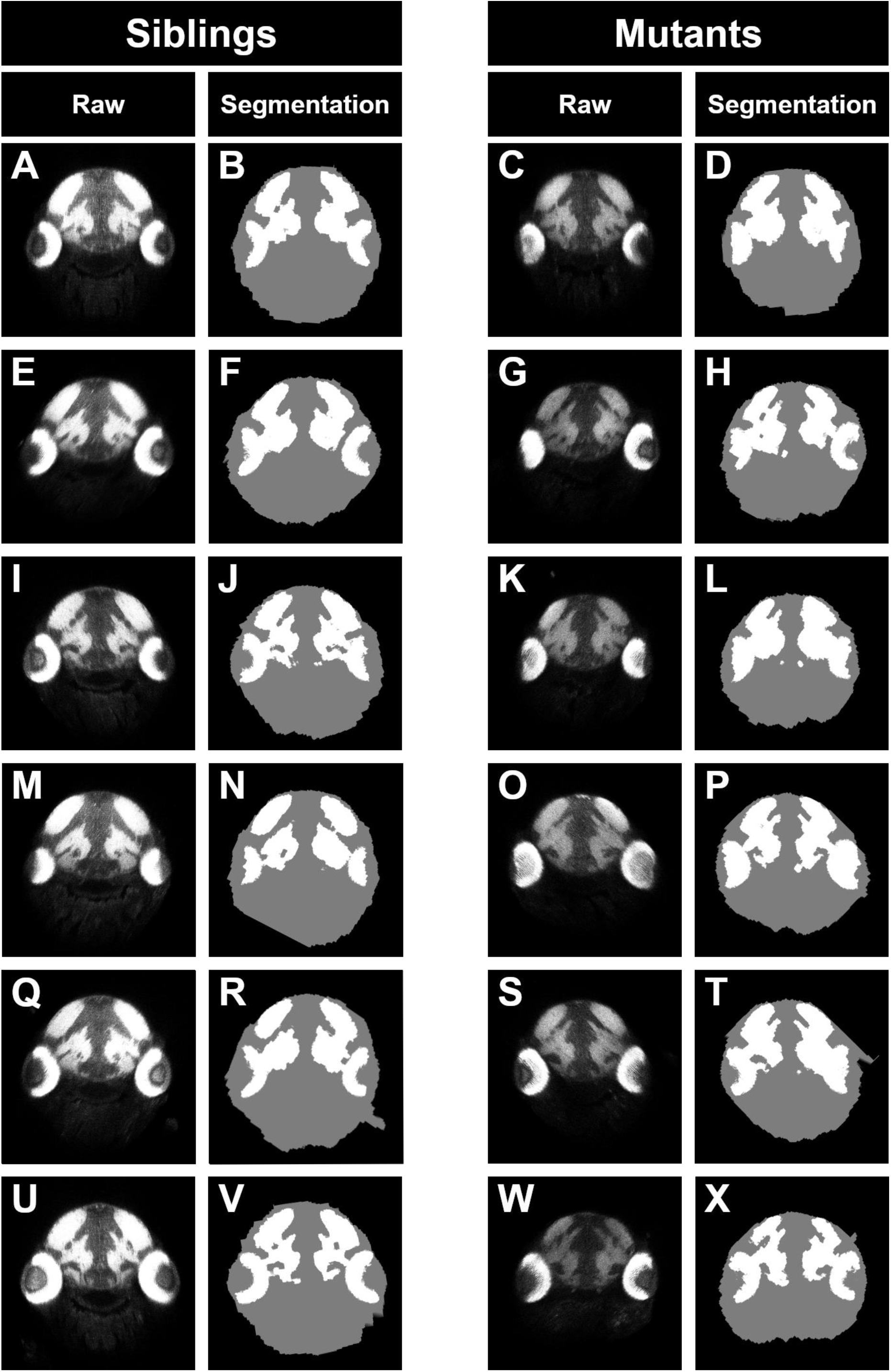
Use of the ZebraFishInspector (ZBI) platform to study midbrain morphology in *wdr12* mutants. The 12 samples shown highlight the homogeneity of brain size in mutants and wild-type EEs. A slight hypomorphy of the whole body (gray disk) and brain white matter (in white) can be observed in the mutants (C, G, K, O, S, W) relative to their siblings (A, E, I, M, Q, U). Nevertheless the quantitative results of the volumetric analysis of 3D images (Fig. 11) reveal much more striking and statistically significant differences in the size of the body and brain white matter between mutants and siblings, demonstrating the advantage of using 3D images. Volumetric analysis, which can be used as the basis of statistical tests, is performed on segmentation results (B, D, F, H, J, L, N, P, R, T, V, X). The accuracy of the segmentation algorithm is demonstrated in this figure.

We then launched the ZBI algorithms on 11 Wt/Htz EEs (one lost sample) and 12 Hmz mutant EEs, to estimate EE and brain white matter volumes. This volumetric analysis clearly revealed that the mutants were smaller than their Wt/Htz siblings (Fig. 10G; Welch test: *p*=5 x 10-08), confirming the overall growth deficits observed in these mutants. Furthermore, despite the much smaller white matter volume in the mutants (Welch test, *p*=3 x 10-06) (Fig. 10H), the volume of the white matter relative to global EE volume was no smaller in the mutants than in the wild type (Fig. 10I), strongly suggesting that this mutation affected brain growth, but that the decrease in brain white matter volume was no greater than the decrease in the growth of other organs, instead being associated with general dwarfism. A visual analysis of the images recorded revealed a normal morphology of the brain.

**Fig. 10.**
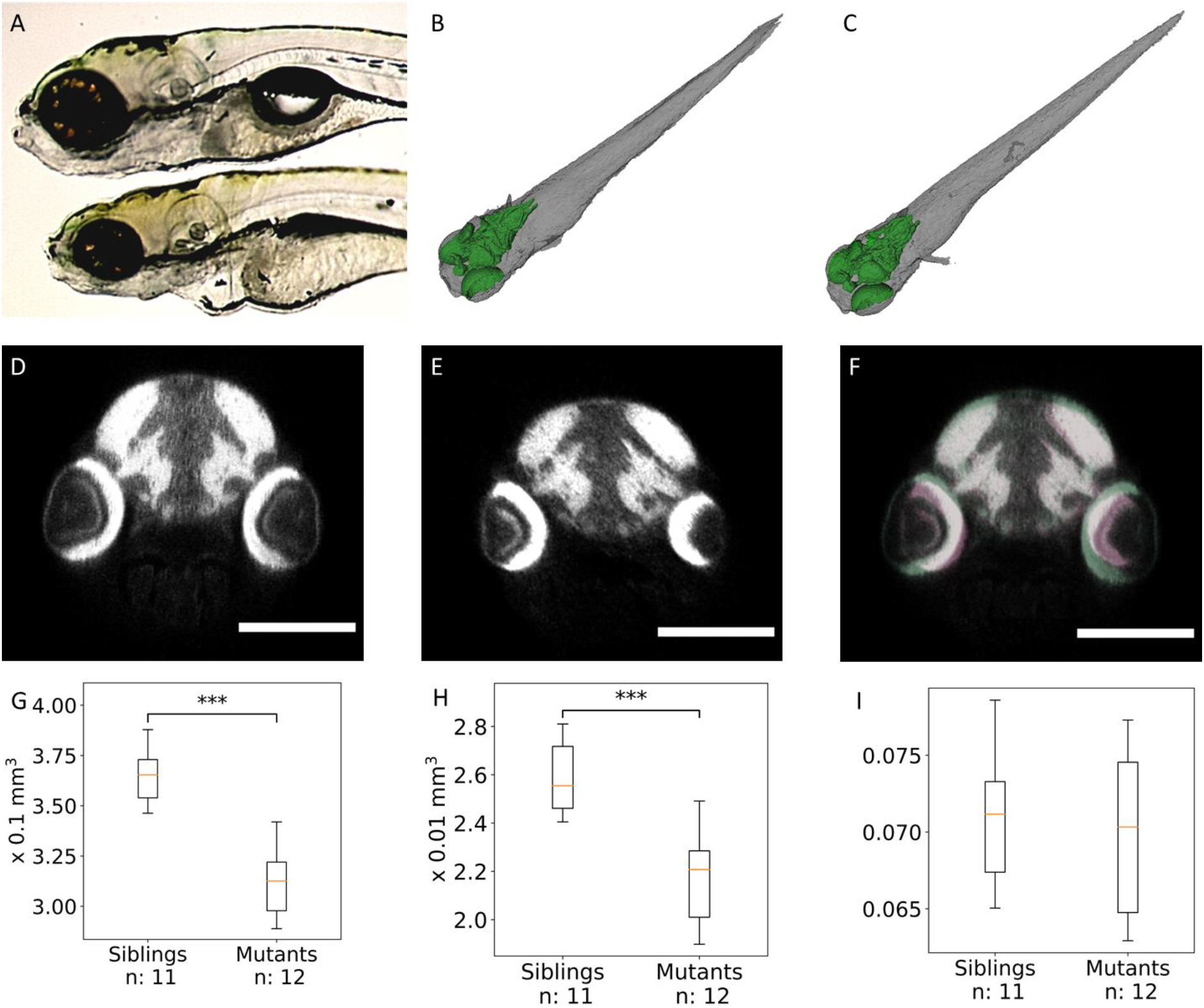
Volumetric analysis of white matter and of the whole body in Wdr12 mutants reveals statistically significant global hypomorphy. (A) Wide-field microscopy of 6 dpf EEs. The wild type is shown at the top, and the *wdr12*^hi3120Tg/hi3120Tg^ mutants below. The mutants have smaller eyes, but it is not clear whether brain size differs between the mutants and their siblings (source Zfin). (B-C) 3D surface rendering of segmentations. EE in gray, white matter in green. These visualizations reveal no obvious difference between mutants and their siblings. (B) WT or Htz mutants (C) Hmz mutants. (D-E) Transverse sections. The mutants clearly have smaller eyes, but it is not possible to distinguish between the brain phenotypes of the mutants and their siblings on these 2D pictures. (F) Superimposition of the transverse sections shown in D and E. Siblings in green, mutants in pink. (G-I) Boxplots of whole EE volumes(G), brain volumes (H) and brain-to-EE volume ratios (I). The mutants are significantly smaller than their wild-type siblings. Brain volume is also significantly smaller in the mutants. However, EE and brain volume decreases are homothetic.

This study highlights the power of our approach for detecting and confirming, with as few as 12 samples, discreet differences, with significantly EE and brain volumes in the mutants, not otherwise clearly visible and which have not been reported before. Fast volumetric analysis successfully generated statistically significant results from as few as 12 3D images (Fig. 10G-H). The identification of brain volume defects in eight mutants is therefore possible by imaging only two dishes containing 48 samples. This proposed method paves the way for other refined analyses involving other brain markers or other organs.

### HCA for assessing the effect of early developmental exposure to environmentally relevant concentrations of 5-fluorouracil

We estimated the performance of the ZeBraInspector platform for large-scale HCI in toxicology, by treating zebrafish EEs with 5-fluorouracyl (5-FU). The active concentrations of 5-FU used were as reported by Kovács et al., (2016), and corresponded to approximately one million times the concentrations found in the environment (Kosjek et al., 2013) and 2 to 40 times the concentrations found in the plasma of human patients treated for colon cancer (Casale et al., 2004).

We tested five concentrations, by placing five batches of 20 recently spawned eggs in six-well plates. We included two controls in each experiment (embryo medium alone, and embryo medium/DMSO). The controls were placed in separate dishes, to prevent any toxicity due to the passage of 5-FU in air. For all treatments, EEs were exposed to 5-FU for five days, starting one hour post-fertilization, with daily renewal of the medium. Each day, mortality was recorded and the dead fish, a maximum of three dead fish per batch (see figure for *n*), were removed, with no significant difference between control and treated EEs (not shown). Most samplesappeared indistinguishable from the controls under the dissecting microscope. Only about 20% of the EEs exposed to 2 g/L 5-FU presented obvious abnormalities: a short body axis and head, microphthalmia, cardiac edema, an abnormal swim bladder and short yolk vesicle (Fig. 11B).

**Fig. 11.**
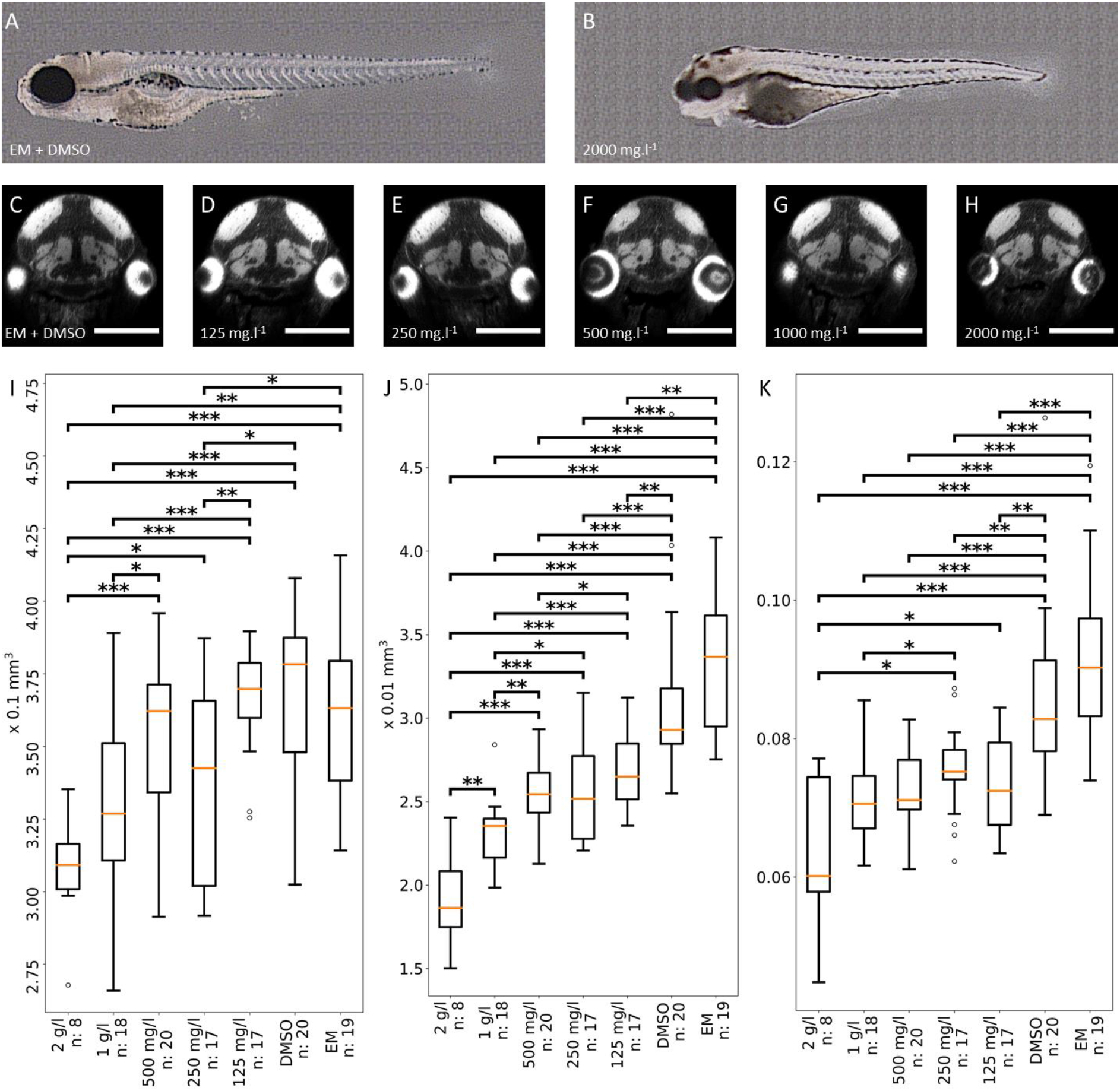
Medically relevant concentrations of 5-FU lead to dwarfism and microcephaly. (A) Dissecting microscope side view of a control EE. (B) Corresponding side view of an EE treated with 2 g/L 5-FU, with a short body axis, cardiac edema, and diverse abnormalities. (C-H) Transverse 2D sections through the midbrains of control zebrafish or fish exposed to 5-FU appear to be similar. Identical levels were selected for brain sectioning. However, due to variations in the orientations of sections, the eyes were not always sectioned at the same level. (AC control EE exposed to DMSO. (D-H) EE exposed to 125, 250, 500, 1000 and 2000 mg/L 5-FU, respectively. Scale bar = 250 μm(G-L) (I-K) Boxplots of whole EE volumes. (I), brain volumes (J) and brain-to-EE volume ratios (K). EEs treated with high concentrations of 5-FU are significantly smaller than controls. Their brains are also significantly smaller. Moreover, white matter/whole EE volume ratios indicate that brains the decrease in brain size is greater than the decrease in overall EE body size, suggesting that 5-FU may specifically affect neurogenesis, causing strong microcephaly.

We measured EE volume with ZBI (Fig. 11I). The EEs incubated with the highest 5-FU concentrations were significantly smaller. This finding contrasts with previous reports that, surprisingly, found no effect or even an increase in fish size due to 5-FU treatments (Kovács et al., 2016). We suggest that these contradictory results may reflect differences in the timing of treatment. Ng et al. (2020) initiated treatment at 5dpf, whereas we initiated treatment less than one hour after spawning, as we hypothesized that this would maximize 5-FU entry into the chorion. Indeed the perivitelline space continues to grow until 1 hpf, due to osmotic water inflow, with the chorion subsequently hardening and becoming less permeable. This hypothesis was confirmed by experiments involving treatment initiation at later time points (>6 hpf), for which no defects were induced, even at high doses (data not shown).

Our analysis revealed a significant decrease in the volume of the brain white matter (Fig. 11J), more marked than that for whole EE volume (Fig. 11I), as shown by the ratio of the white matter and EE volumes (Fig. 11K). Interestingly, this difference was not obvious on transverse sections (Fig. 11C-H).

Thus, using 3D images, we were able to detect discreet defects that were not otherwise obvious.

Overall, these results from our platform reveal that the exposure of zebrafish embryos to medically relevant concentrations of 5-FU very early in growth affects growth and brain white-matter size, as expected for treatment with an antiproliferative agent.

## Conclusion and Perspectives

We have developed a fast and flexible high-resolution 3D image acquisition and analysis platform for HCI on 5 dpf EEs: ZeBraInspector. The basic volumetric analysis of this pipeline can be performed with simple lipophilic dye labeling, but more specialized phenotypes can be investigated by integrating immunohistochemical labeling for appropriate epitopes or using transgenic lines expressing a fluorophore in the region/tissue of interest. Standardized rearing, staining, mounting and imaging techniques ensure consistent quality and the obtainment of the large number of 3D images required for statistical analysis at population level. As a convenient way of interacting with the automated volumetric analysis, a graphical user interface enables users to review data quality and to sample the population on the basis of preprocessed, lightweight representations of the raw data. This interface also can be used for the analysis of highly variable phenotypes resulting from zebrafish mutants. For example, it can be used to sort samples with various degrees of defect and to analyze transgene expression or antibody labeling patterns more reliably.

The modular nature of our processing pipeline makes it possible to integrate additional analysis modules, which can be developed specifically for a given application. The active development of additional analysis modules is currently focused on the integration of machine learning, to accelerate the image segmentation process for volumetry, and non-rigid registration to increase the sensitivity of detection forvolumetrically neutral deformations within the sample. For larger studies, we foresee an increase in productivity through the further robotization of specimen handling during mounting and image acquisition, through the use of 96-well plates and, if possible in the study concerned, a decrease in sampling rate.

This pipeline was tested in studies of subtle volumetric aberrations in transgenic mutants and of the neurodevelopmental impact of an ecotoxic compound. It could be modified and scaled to any application involving investigations of the impact of test compounds on an entire, physiologically intact vertebrate model organism, at a holistic level.

## Supporting information

Guidelines for ZBI

## Acknowledgements

We wish to thank Marie-Elise Schwartz and Julien Hémon for skillful fish rearing. We thank Jean-Pierre Levraud for sharing pictures of two transgenic lines, and Koji Ando for the kdlr-DsRED line. We thank Elodie De Job and Laurie Rivière for their expert technical contributions to the development of our tissue-clearing procedures, Anna Renoux for the genotyping of *wdr12* fish, Maxence Frétaud, Christelle Langevin and Violette Thermes for their contributions to protocol development through helpful discussions in the framework of the TEFOR phenotyping group. We also thank Lionel Moreira and Vincent Jourdain for their contribution to the TEFOR portfolio webpage, and Giovanni Cherchia and Barbara Rizzi for advice concerning algorithm developments. Kleio Petratou, Ali Kiai provided helpful comments on the ZBI software. We thank Johanna Djian-Zaouche for discussions and support. This work was supported by ANR-TEFOR-”Investissement d’avenir” (ANR-II-INBS-0014), ANR Fish-RNAvax (ANR-16-CE20-0002-01), ANR FEATS (ANR-19-CE34-0005-05) and institutional grants from the CNRS and INRAE. We warmly thank the Leducq foundation for the RETP grant and constant support, which was central to the development of the ZBI platform. SL received a PhD grant from Université Paris-Est.

