## Supplementary material for "ZeBraInspector, a platform for the automated segmentation and analysis of body and brain volumes in whole 5 dpf zebrafish following simultaneous visualization with identical orientations": Guidelines for ZBI

### Supplemental data

#### Guidelines for high-content images (HCI)

In our pipeline, the software of our microscope generates image stacks by merging individual tiles. However, in other contexts, stitching procedures may be performed by the user before image processing. It is recommended to perform stitching by first reconstructing the volumes from each image tile and then performing 3D stitching.

Different image sizes can be processed by our algorithm. Image processing is performed with the voxel as a standard unit of measurement. The use of isotropic voxels is recommended.

We encourage the recording of images with a 12-bit depth to obtain a good representation of the dynamic range of the digitized datasets. A correctly acquired image should have an equilibrated grayscale value histogram. All the grayscale values allowed by the 12-bit encoding should be used, while limiting saturation. If the image uses only a portion of the grayscale value range, histogram normalization may be used.

It is also important to maximize the contrast within the image. For dye acquisition, the laser power should be chosen so as to saturate only the photoreceptor layer of the retina, limiting the saturation of other tissues to a minimum. It is also preferable to keep the background noise outside the sample as low as possible. The choice of offset should, thus, limit the number of non-zero voxels outside the sample.

Images must be saved in a format allowing the preservation of metadata. The nd2, tiff and mha formats are capable of preserving this information and are compatible with the ZBI software. For mha and tif files, each channel must be stored as a separate image, and the file name should end with _C00, _C01 or _C02. Sample tracking through the different phases of the experiment is a major goal requiring a solid data management plan. As a means of maintaining flexibility in the choice of analytical tools and accessibility to diverse users, we developed a data management system based entirely on structured file names (Materials and Methods).

### Guidelines for the ZeBraInspector (ZBI) software

Help can be found by clicking on **?** on each screen of the software.

**Download the software** from https://tefor.net/portfolio/zebrainspector, execute it (a security window may appear; click on complementary information and execute). It may take one minute to launch the software.

**The software was developed for 5-day zebrafish EEs.** With time, we will validate its use for other stages and other species. We will also try to expand its use to other microscopes. Please keep us informed of your results with other types of tools and samples at!

If needed, you can also **download images for demos** from https://tefor.net/portfolio/zebrainspector

**Screen resolution:** The software was optimized for use with 1920x1080 screens. Large screens, of course, improve rendering.

**Image format and depth:** Only nd2, mha and tif formats are accepted. Image analysis algorithms were developed on 12-bit unsigned integer-encoded images. These algorithms do not work with lower-level encoding, as for 8-bit images. However, they may work correctly at higher levels of integer encoding, for 16-bit images, for example.

**Image specifications:** Whole EEs are required for registration algorithms, and it is better to have the head on the left of the image. It is recommended to use a computer with at least 32 Go of RAM that will run smoothly with 2500x1000x500 voxel images (voxel size about two microns, voxels must be isotropic). Objectives with a field of 0.5 millimeters: 12x2 tiles. Objectives with a field of 1 millimeter: 6x1 tiles. Performance increases with RAM: larger images and/or larger numbers of images can be loaded in the software.

**Configuring the interface:** The numbers of lines and columns to be displayed per screen and of channels to be viewed. At the right of the top menu bar, select zoom mode: single image, lines, rows or all visible images on the screen. You can zoom by clicking on an image.

**The contrast** of the reference dye channel must be precisely tuned. No specific parameter is needed (more guidelines are provided in the microscopy chapter above). With depigmented samples, saturate the retina layers slightly and check that the lens and background have grayscale values of zero. For samples with pigmented eyes, check that the membrane over the eye is clearly visible, to as great a depth as possible in the sample.

**Downsampling factor.** When a file of 2500x1000x500 voxels is loaded, its resolution is downsampled four times by default. The dynamic range for rendering is reduced to 8 bits. Hence, the software loads 100 files of this type into memory with only 2.4 Go of RAM requested. Avoiding downsampling (left in the top menu bar) results in better rendering but may be slow for large numbers of samples. Adapt to achieve the best compromise between sample number, size and rendering quality.

**Optical sections.** The H, S and T buttons in the top menu bar display horizontal, sagittal and transverse views, respectively. Moreover, a rotation around the anteroposterior axis can be performed. Visualizations are maximum-intensity projections of thick optical sections of the sample around the chosen slice. Section size can be selected within the range 4 to 128 µm.

**Snapshot capture.** The observation of a particular structure can be facilitated, or observation can be optimized before capture, by tuning the contrast by clicking on CH O 1 2 in the bottom menu bar. Click on the snapshot button below each image. Image extraction may take one minute. Then save highresolution versions of selected images in a folder (the original file name is changed, the sample numbers in the current session are retained, capture parameters and dates are retained in the file name).

**Segmentation**

Segmentation cannot be performed on registered data because this computation involves changes in pixel intensity along the *xz* axis, which is reoriented by the registration step. To start segmentation, a reference/dye channel and output channel must be selected. Further computations are performed on raw data for the dye/reference channel and the results are displayed in the output channel. If the output channel is populated, the segmentation will replace the previous image.

By default, results are saved in mha format. A checkbox is provided to allow the user to opt for the tiff format rather than the mha format. Before processing, the software asks for a destination folder. Segmentation results are saved in the chosen folder.

Segmentation performed on a 1 Gb image takes about 10 minutes, depending on the computer hardware.

**Registration**

To start registration, select a reference/dye channel. As for segmentation, further computations will beare performed on raw data from the chosen channel. If the registration of segmentations is required, it should be performed just after registration or the segmentation output files should be reloaded with the initial raw data.

For registration, three options are proposed: first, the software can run with $H_2O_2$-depigmented samples or on mutants lacking eye pigmentation. Segmentations are performed with the lenses, which appear as black spheres because they absorb light. Other mutants (such as casper or nacre) have no body pigmentation, but nevertheless have pigments in the eyes. In this case, the depigmentation step is not performed, to preserve native fluorescence, for example. A second “pigmented eyes” (no $H_2O_2$ treatment) option is proposed for these samples. The registration algorithm uses eye shape in this case. A third registration does not use the eyes at all and instead bases the calculation on an ellipsoid bounding the white matter signal, using the axes of this ellipsoid for alignment. This approach may result in less precisely reoriented samples.

As with the segmentation algorithm, a checkbox is provided to allow users to opt for output in the tiff format rather than in the mha format, and the software asks for a destination folder before computation.

For an image containing three channels of 1 Gb, computation may take five minutes per sample, depending on computer hardware.

**Volumetric analysis results in an Excel table** A lipophilic dye (DiI ,Dio) stains the myelin sheaths of neuron fibers, which comprise a domain of the nervous system known as white matter. At 5dpf, this staining provides a good overview of brain morphology. EE volume is the number of pixels contained in the segmented EE, providing a measurement of its size (many other measures may be added).

To start volumetric analysis, select images with correct segmentations by checking the “selected” checkbox below each image. Click on the volumetry button in the bottom menu bar. During the volumetric analysis, the numbers of voxels in each segmentation are counted. As voxels are 3D elements, these numbers can be extrapolated to volumes. A third value, the ratio of white matter volume to the volume of the whole sample, is also calculated. These three values are stored in a spreadsheet for further statistical analysis.

### Troubleshooting

**Software does not start**

ZBI was developed to work in the Windows or Linux environments. A binary file is available for Windows. For the Linux environment, codes are available from a gitlab repository, and the software can be compiled locally. We did not test the running of ZBI codes in Mac environments, and cannot ensure the correct running of the software in such environments. The main window of ZBI should be displayed within about one minute. If the software does not start, please check that the computer has a C++ compiler. For Windows environments, visual C++ 2017 should be installed.

**Software freezes**

This software was designed to work with data of 1 Go per channel, on a computer with at least 32 Gb. It may not be possible to treat the data with smaller RAM dimensions, resulting in software freezes.

Closing the registration or the segmentation windows will not stop the processes from running. Please wait for the end of one computation before starting another, to prevent RAM overload and software freeze. Navigation within the software is slower during computations.

**Failure of registration**

In this case, the image is probably not well-managed by the algorithm due to a low signal/noise ratio for the dye signal, or saturation of the dye signal in domains that are too large. Insufficient sample clearing leading to signal loss at depth may also prevent registration (see Lempereur et al., 2020 in MMTA for a contrast improvement algorithm for confocal images). Please check protocols and guidelines for image acquisition. Obviously, registrations in the casper background are often less successful than those based on lenses with depigmented eyes, because eyes are not necessarily of the same size, are not spherical and eye boundaries are hardly visible ventrally because they are hidden by the more dorsally located eye pigments during imaging.

Also, strongly deformed EEs may not be correctly recognized by the software.
